# Early-life Oxytocin Rescues Hippocampal Synaptic Plasticity and Episodic Memory in a Mouse Model of Fragile X Syndrome

**DOI:** 10.1101/2024.12.20.629802

**Authors:** Jasmine Chavez, Aliza A. Le, Julie C. Lauterborn, Brittney M. Cox, Yousheng Jia, Gary Lynch, Christine M. Gall

## Abstract

Cognitive disabilities including impairments to episodic memory are debilitating features of autism spectrum disorder (ASD). Here we report that early-life treatment with oxytocin (OXT) fully restores episodic memory and associated synaptic plasticity in a rodent model of an ASD. *Fmr1*-knockout (KO) mice -- which mimic the single gene mutation in Fragile X Syndrome -- failed to encode three basic elements of an episode (identity, location, and temporal order) during a first time, unrewarded encounter with a set of cues. Intranasal administration of OXT during the second postnatal week eliminated each of these impairments in mice tested in adulthood. OXT treatment during the second, but not fifth, postnatal week also corrected pronounced defects in two distinct forms of hippocampal Long-Term Potentiation (LTP). Rescue of LTP in lateral perforant path (LPP) to dentate gyrus synapses was linked to recovery of NMDAR-gated synaptic responses, which are otherwise profoundly reduced in the mutant LPP. LTP induced by a threshold theta burst stimulation protocol in CA3-CA1 synapses was severely impaired in adult *Fmr1*-KOs as was a previously unreported post-induction growth phase for potentiation. Both effects were restored in adult Fmr1-KOs given early OXT treatment. Infusion of OXT into adult *Fmr1*-KO hippocampal slices normalized LTP in CA1 but had no effect on the defective potentiation (or NMDAR-mediated EPSCs) at LPP-dentate gyrus synapses. These results show that severe, autism-related defects in cognition critical to memory, as evident in a rodent model, are reversible and that an early-life therapeutic intervention can effect an enduring restoration of function.

## INTRODUCTION

Fragile X syndrome (FXS) is the most common inherited form of intellectual disability and has a high comorbidity for autism spectrum disorders (ASDs)^1^. In addition to learning impairments, the FXS phenotype includes sensory hyperexcitability, seizures, anxiety, and sociability deficits^2–4^. FXS arises from a trinucleotide repeat expansion in the 5’ untranslated region of the *Fmr1* gene leading to its hypermethylation and silencing^5–7^. Pharmacological interventions for FXS individuals have primarily focused on symptomatic treatment of behavioral and psychiatric disturbances (e.g., anxiety, ADHD, mood instability and aggression)^8,9^. Notably, there has been little progress in developing treatments for the memory and cognitive problems that accompany the disorder.

Recent studies have investigated the possibility of using the hypothalamic neuropeptide oxytocin (OXT) to treat certain psychiatric disturbances associated with autism^10–12^. OXT plays an important role in social behavior^13,14^, and reduced plasma levels of the hormone have been observed in autistic individuals^10,15,16^. OXT expression is also depressed in the hypothalamic paraventricular nucleus (PVN) of *Fmr1*-knockout (KO) mice^17,18^. These findings suggest that deficits in OXT production could contribute to sociability issues associated with FXS and other congenital ASDs. In support of this, Peñagarikano et al. found that administering intranasal OXT daily from postnatal day (P) 7 to P21 increased PVN OXT levels and eliminated social behavior disturbances in the *Cntnap*2-KO mouse model of autism^19^. Other work has corroborated this early-life effect of OXT and showed that a narrower treatment window (P12-P16) improved the social behavior of *Fmr1*-KOs as evident in tests conducted at 2 months of age^17^. Whether the beneficial effects of early-life OXT treatments extend beyond sociability to cognitive problems associated with autism-related disorders has not been examined.

Multiple studies have found that episodic memory, a form of everyday encoding that is critical for orderly thinking^20,21^, is impaired in autistic individuals^22–24^. Episodic processing organizes the flow of everyday experience into autobiographical units that include context-dependent information about the identities (‘what’), locations (‘where’), and serial order (‘when’) for a set of cues. This is accomplished without practice or overt rewards. Acquisition and retrieval of episodic information is heavily dependent upon the hippocampus^25,26^, a structure that is significantly disturbed in *Fmr1-*KOs. In particular, while baseline synaptic transmission appears to be normal in mutants, two distinct forms of memory-related long term potentiation (LTP) are substantially reduced in magnitude^27–30^. Defects in hippocampal plasticity could thus be major contributors to autism-related episodic memory problems. The hippocampus is innervated by OXT-positive fibers originating in the PVN^31^ and expresses moderate levels of OXT receptors^32,33^. Accordingly, we tested the effects of early-life OXT treatment on both LTP and episodic memory in adult *Fmr1*-KOs. The results revealed an enduring positive effect of OXT treatment in the mutant indicating that early-life interventions may be feasible for the broader phenotype of FXS.

## METHODS

### Animals

Studies used male FVB129 *Fmr1*-KO and wild type (WT) mice from an in-house colony. After weaning at P21, mice were group-housed with same sex littermates in rooms at 68°F and 55% humidity, 12 hr on/12 hr off light cycle. Food and water were provided *ad libitum*. Protocols were approved by Institutional Animal Care and Use Committee at University of California, Irvine and are consistent with the NIH Guide for the Care and Use of Laboratory Animals.

### Intranasal Oxytocin Treatment^19^

Oxytocin (1 µg/µL) or saline (SAL) was administered intranasally (2 µL per nostril) once daily for 7 days between P7-P13 or P30-P36. Human OXT (CRO300A, CellSciences) was kept as a 10 mg/mL stock and diluted in 0.9% NaCl for administration. Intranasal OXT and SAL treatments were done in parallel for a given cohort. After the last treatment, the pups were left undisturbed with their dam and littermates until weaning. Behavioral and electrophysiological experimentation began at 2-4 months of age.

### Behavioral Assays

Behavioral sessions were performed between 10AM-3PM and recorded using an overhead camera; videos were scored by 2 observers blind to group. Each mouse was used for at most two paradigms with 3 days between tasks. All experimental chambers had distinct visual cues on the walls. The chambers, objects and scent jars were cleaned between uses with 70% ethanol. Animals were excluded from analyses if they sampled any cue for <1s during the exposure trial, excluding the episodic ‘Where’ task. In all cases, the time the animals had incidental contact, gnawed, or climbed on the object/cup were excluded from analyses. statistical analyses of discrimination indices and raw sampling times for each behavior are presented in supplementary **Tables S2** and **S3**, respectively.

#### Object Location Memory (OLM)^28,34–36^

Mice were habituated to an empty 30×25-cm (21.5-cm height) plexiglass chamber (5-min/day) for 2 days. The exposure trial consisted of 5 min exploration of the chamber containing two identical glass funnels positioned in two adjacent corners. For the test trial 24h later, the mouse was allowed 5 min to explore the chamber with one funnel displaced toward the center chamber. The time (t) the animal spent sampling the objects (nose oriented towards and within 0.5-cm of the object) was quantified in seconds. Performance was assessed using a discrimination index (DI): 100 x [(t_novel_ − t_familiar_) / t_total_]. The distance traveled was measured using EthoVision XT (Noldus).

#### Three-Chamber Sociability Task^28,37^

For habituation to the 3-chamber plexiglass arena (63×41-cm floor, 25-cm height) the test mouse was allowed to explore the empty center (start) chamber for 5 min, all three chambers for 10 min, and then the center chamber for 5 min. <u>Approach</u>: Immediately after habituation, the animal was allowed 10 min to explore the three chambers with one side chamber containing an empty inverted cup, and the other containing a ‘stranger’ mouse (S1) inside an inverted cup; stranger mice were unfamiliar age-matched FVB129 males. The test animal then rested in the center chamber (5 min) while S1 was moved to the chamber previously containing the empty cup, and a second stranger mouse (S2) was placed in an inverted cup in S1’s previous location. <u>Recognition</u>: The test animal was allowed 10 min to explore the 3 chambers. The time the test mouse oriented towards and within 0.5-cm of a mouse or empty cup was scored. For social approach, DI=100 x [(t_mouse_ – t_object_)/ t_total_]. For social recognition, DI = 100 x [(t_stranger_ – t_familiar_) / t_total_].

#### Two-odor Discrimination and Episodic Memory Tasks^29,38^

These tasks tested for detection of a specific novelty using odorant cues at a concentration of equal salience for WT and *Fmr1*-KO mice^39^. All odorants (see supplementary **Table S1**) were dissolved in mineral oil to ∼0.1 Pa. and pipetted (∼100-120 µL) onto filter papers placed inside glass jars (5.2-cm diameter, 5-cm tall) sealed with perforated lids. All tasks utilized the OLM arenas except the ‘Where’ task which employed a 60×60-cm (30-cm height) plexiglass arena; all floor of the arena had shallow indentions to maintain jar placement across trials. In all cases, mice were first habituated to the arena containing unscented jars (5 min). The jars were then removed and after a 5-min delay, the animals were tested in the following paradigms and DIs were calculated as described.

#### Simple 2-Odor Discrimination^29,38^

The mouse was presented with an identical odor pair (E:E) for 3 min and then the scented jars were removed. After 21 minutes (to match the time elapsed from initial odor exposure to testing in the Serial Odor ‘What’ task), the animals were exposed to a new odor pair including familiar odor E and novel odor F (E:F). For the 2-odor and Serial ‘What’ (below) paradigms, the DI was calculated as: 100 x [(t_novel_ – t_familiar_)/ t_total_].

#### Episodic Serial Odor ‘What’^29,38^

The mouse was presented with a series of three identical odor pairs (A:A, B:B, C:C) for 3 minutes each, with 5 min spacing between pairs. Five minutes later, the animal was presented with two test odors including the first odor in the sample sequence (A) and novel odor D (A:D).

#### Episodic Serial Odor ‘When’^38^

Initial odor sampling was the same as in the ‘What’ task, but in this paradigm, the animal sampled a series of four sample pairs (A:A, B:B, C:C, D:D). For the test trial, the mouse was presented with an odor pair including less recently experienced odor ‘B’ and more recently experienced odor ‘C’. DI= 100 x [(t_older_ – t_recent_)/ t_total_].

#### Episodic 4-Corner ‘Where’^38^

Following a 5 min habituation session with four odorless cups placed near the arena corners, the empty cups were replaced with cups each containing a distinct odor (A, B, C and D). After 5 min of exploration, the animals were returned to their home-cages. 24-hrs later, the animals were allowed 5 min to explore the arena containing clean cups scented with the initial 4 odors but with the positions of two diagonally opposite odors swapped. DI= 100 x [(t_mean of switched pair_ – t_mean of stationary pair_)/ t_total_].

### Extracellular Field Electrophysiology

Acute hippocampal slices from adult mice were prepared for analysis of LTP of the CA3-CA1 Schaffer-commissural (S-C) afferents to CA1 and lateral perforant path (LPP) as described^29,35,36^. For S-C recordings, slices were collected into chilled oxygenated artificial cerebrospinal fluid (aCSF)^40^. For LPP recordings, slices were collected into ice-cold high magnesium-aCSF^40^. For both S-C and LPP preparations, slices were immediately placed onto an interface recording chamber with continuously perfused aCSF (31±1°C). Experiments were initiated ∼1.5-2 hrs later. Baseline responses were collected for 20 min before inducing LTP (see below) and for 1 hr thereafter.

#### S-C LTP

The stimulating electrode was placed in CA1c stratum radiatum (SR), and the recording electrode was placed in CA1b SR, both equidistant from the CA1 stratum pyramidale^36^. LTP was induced by applying a three burst train of theta burst stimulation (TBS: four pulses, 100 Hz; 200 ms between bursts) at baseline stimulation intensity.

#### LPP-LTP

Stimulating and recording electrodes were placed in the dentate gyrus (DG) outer molecular layer, and stimulation of the LPP afferents was confirmed by the presence of paired pulse facilitation with a 40-ms interpulse interval^41,42^. LTP was induced using a single train of high-frequency stimulation (HFS: 1 second, 100 Hz) at 2x duration and 1.5x intensity of baseline stimulation^29^.

Initial slopes and amplitudes of field excitatory postsynaptic potentials (fEPSPs) were digitized at 20kHz using an AC amplifier (A-M Systems, 1700) and recorded using NACGather 2.0 (Theta Burst Corp., Irvine, CA). The magnitude of LTP was assessed by comparing the mean of the fEPSP slopes collected 55-60 minutes after TBS or HFS to the mean baseline response.

### Whole-Cell Current-Clamp

Hippocampal slices were prepared as described^40,43^ and placed in a submerged recording chamber continuously perfused at 2–3 mL/min with oxygenated aCSF at 32°C. Whole-cell recordings were obtained from DG granule cells with 4–7 MΩ recording pipettes filled with a cesium-based solution^44^. The stimulating electrode, placed in the DG outer molecular layer, was used to evoke LPP-specific responses. Excitatory postsynaptic currents (EPSCs) were recorded by clamping the granule cell at −70 mV in the presence of 50-µM picrotoxin^40^. NMDAR-to AMPA receptor (AMPAR)-mediated EPSC ratios were calculated as the ratio of the current at 30 ms after stimulus onset with −10 mV holding potential to the peak current at −70 mV^43^. All recordings were analyzed offline using Clampfit software (Molecular Devices, San Jose, CA).

#### Oxytocin treatment of hippocampal slices

After collecting 20 min stable baseline responses, OXT (10 µM in water) or vehicle was infused into the slice chamber using a syringe pump (6mL/hr; KD Scientific model 100; the bath concentration of OXT was 1 µM.

#### Statistical analysis

Significance was determined using GraphPad Prism v6.0. Two-way ANOVA with Tukey post-hoc analyses was used for genotype and drug comparisons. Frequency facilitation plots and TBS responses were analyzed with repeated-measures two-way ANOVA. Linear regression of slope was used to assess input/output curves.

## RESULTS

To assess the effects of early-life postnatal OXT treatment on synaptic plasticity and memory in adulthood, *Fmr1*-KO and WT mice were given daily intranasal OXT or isotonic SAL treatments for one week (P7-P13). Testing commenced when mice reached 2 months of age. The majority of animals were evaluated in one or two behavioral tests prior to electrophysiological studies (**Fig. 1A**).

**Figure 1:**
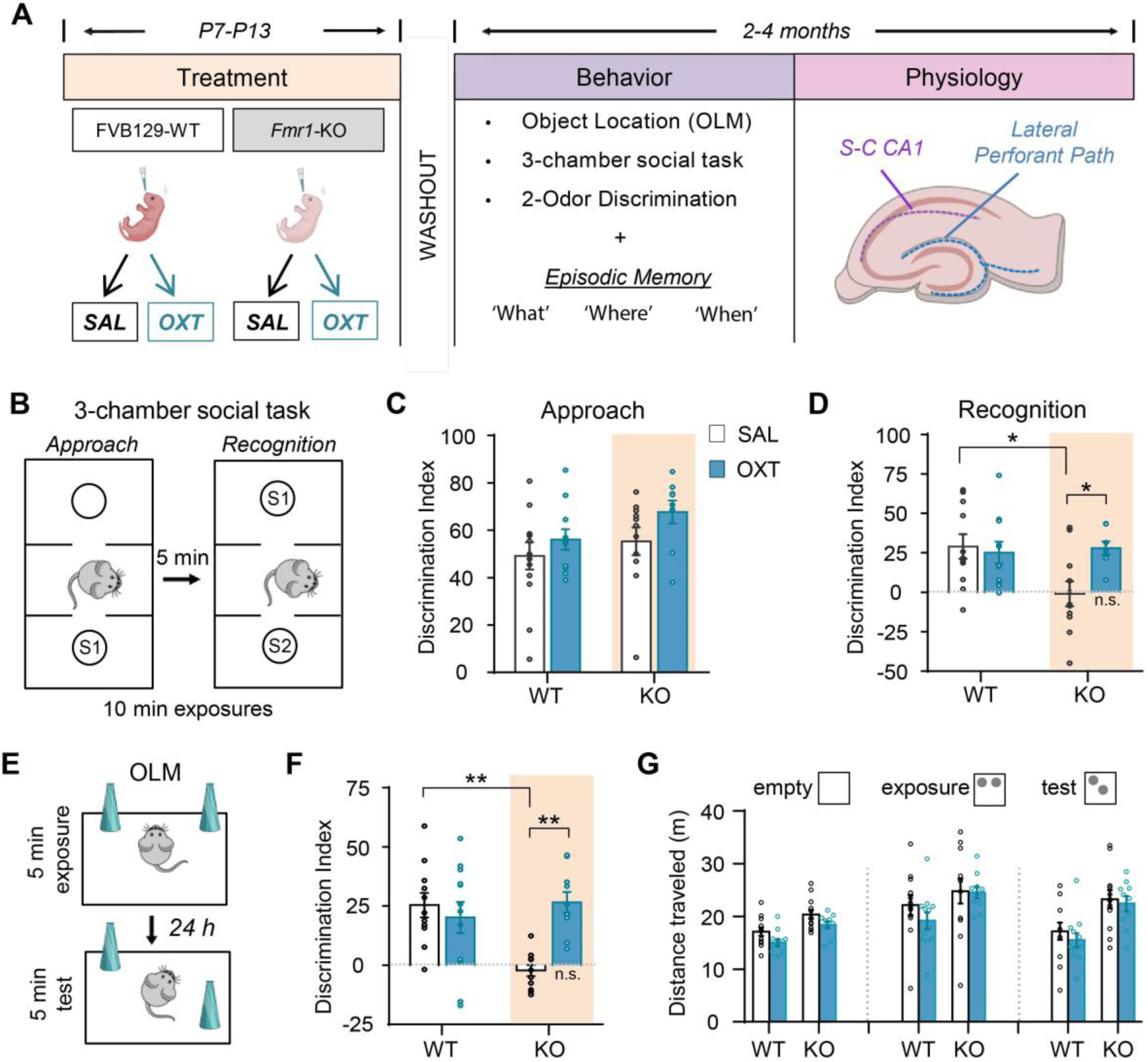
Postnatal oxytocin (OXT) treatment improves social and spatial memory in adult *Fmr1*-KO mice. (**A**) Schematic of timelines for intranasal OXT and saline (SAL) treatment in *Fmr1*-KO (KO) and wild-type (WT) mice (daily from postnatal day (P) 7 to P13), and testing for behavioral and hippocampal electrophysiological effects of genotype and treatment in 2-4 mo old adults. (**B**) The 3-chamber social task entailed two phases. *Approach*: Mice were exposed to an empty cup or stranger mouse (under cup) (S1). *Recognition*: Mice were allowed to explore the apparatus with the side chambers containing a novel stranger mouse (S2) and the initial stranger mouse (S1). (**C**) In Approach, all groups preferentially explored the stranger mouse over the empty cup as indicated by high discrimination index scores (F_(1, 40)_=0.255, p=0.617, 2-way ANOVA (interaction). (**D**) In Recognition, SAL-treated KOs did not discriminate the stranger mouse (S1 vs S2) (F_(1,37)_=5.149, p=0.029; *p=0.02 KO-SAL vs. WT-SAL), whereas OXT-treated KOs did (*p=0.047 KO-OXT vs. KO-SAL) comparable to behavior of the WTs (n.s. p=0.99 KO-OXT vs. WT-OXT). (**E**) Object location memory (OLM) paradigm. Mice were exposed to two identical objects placed near adjacent corners. After 24 h, the mice were reintroduced to the chamber in which one object was displaced. (**F**) Both WT groups preferentially explored the novel-location object whereas SAL-treated KOs did not (F_(1,40)_=7.90, p=0.008; **p=0.002). *Fmr1*-KOs given OXT preferentially explored the displaced object (**p=0.0015 KO-SAL vs. KO-OXT) with discrimination indices comparable to the WT groups (n.s. p=0.799 KO-OXT vs. WT-OXT). (**G**) Distance traveled during the OLM habituation (empty chamber), exposure, and test sessions. There was a consistent effect of genotype (empty chamber: F_(1,42)_=18.77, p<0.0001, exposure: F_(1,42)_=4.636, p=0.0371, and test: F_(1,42)_=17.52, p=0.0001) in that KO mice travelled more than WTs; post-hoc analysis showed that OXT treatment did not influence the distance traveled by KO or WT mice in any session. *Statistics (C, D, F, G)*: two-way ANOVA (interaction) with Tukey post-hoc comparisons indicated as: *p<0.05 and **p<0.01.

### Early-life OXT treatment rescues social recognition in adult Fmr1-KO mice

Early-life OXT treatment has been shown to normalize social behavior in mouse ASD models^45^ including *Magel2*-deficient^46^, *Cntnap*2-deficient^19^ and *Fmr1*-KO^17^ mice. The latter study demonstrated an enduring rescue of social recognition behavior following twice daily intranasal OXT treatment from P12-P16. We first tested if OXT given once daily from P7 to P13 normalizes social behavior in *Fmr1*-KOs tested as adults. We used the 3-chamber paradigm (**Fig. 1B**) to assess social approach and social recognition^37^. In accord with previous reports^17,28,47^, *Fmr1*-KO and WT mice exhibited comparable social approach, preferentially exploring a mouse (S1) over the object regardless of treatment (**Fig. 1C, S1A**). In the subsequent social recognition trial, both WT groups preferentially explored the novel mouse (S2) over the familiar mouse (S1). This discrimination was absent in *Fmr1*-KOs given SAL but was restored to WT levels by OXT (**Fig. 1D, S1B**). These results corroborate reports that early postnatal OXT treatment rescues social behavior in *Fmr1*-KOs and that this effect persists into adulthood.

### Object Location memory (OLM) is restored by postnatal OXT in Fmr1-KO mice

*Fmr1*-KO mice have deficits in OLM^34^, a task that depends on hippocampus^48,49^. We tested if postnatal OXT treatment (P7-P13) improved OLM (**Fig. 1E**) in adult *Fmr1*-KOs. Wild-types given SAL preferentially explored the displaced object at testing whereas SAL-treated KOs did not, confirming the genotype-associated deficit in spatial memory^28,34^. In contrast, *Fmr1*-KOs given early-life OXT preferentially sampled the displaced object and did so to a similar degree as SAL-treated WTs (**Fig. 1F, S1C**). Oxytocin treatment had no effect on Discrimination Indices (DIs) in WTs (**Fig. 1F**). Analysis of distance traveled during OLM habituation (empty chamber), exposure, and test sessions revealed that *Fmr1*-KO mice ran somewhat more than WTs, and OXT treatment did not influence this difference (**Fig. 1G**).

### Episodic memory in adult Fmr1-KOs is rescued by postnatal OXT

Autistic individuals experience difficulties in processing episodic memory^22,23^ and it is thought that these impairments contribute to the intellectual disabilities associated with FXS and ASD. We previously demonstrated that *Fmr1*-KOs have a pronounced deficit in the unsupervised encoding of cue identity (episodic ‘what’ information)^29^. Here we extended this analysis to effects of genotype on acquisition of ‘where’ and ‘when’ elements of an episode and tested if early OXT treatments correct deficits otherwise evident in adult *Fmr1*-KOs. The protocols did not involve prior training and used odors, which are of inherent interest to rodents, as cues^38^. We first tested if mice from the four groups (WT±OXT; KO±OXT) differed in performance of a 2-odor discrimination task in which they sample odor A and then, after a brief delay, are tested with A vs. novel odor D (**Fig. 2A**). All groups preferentially explored the novel odor, indicating that simple odor discrimination and related memory were unaffected by genotype or OXT treatment (**Fig. 2B, S2A**). Next, we tested mice in the multi-cue, episodic ‘what’ task, in which they were exposed to a sequence of odor pairs (A-A, B-B, C-C), followed by a pair containing familiar odor A and novel odor D (**Fig. 2C**). Both OXT- and SAL-treated WTs preferentially sampled the novel odor whereas SAL-treated KOs did not, consistent with previous findings^29^. In contrast, KOs given OXT performed at WT levels (**Fig. 2D, S2B**).

**Figure 2:**
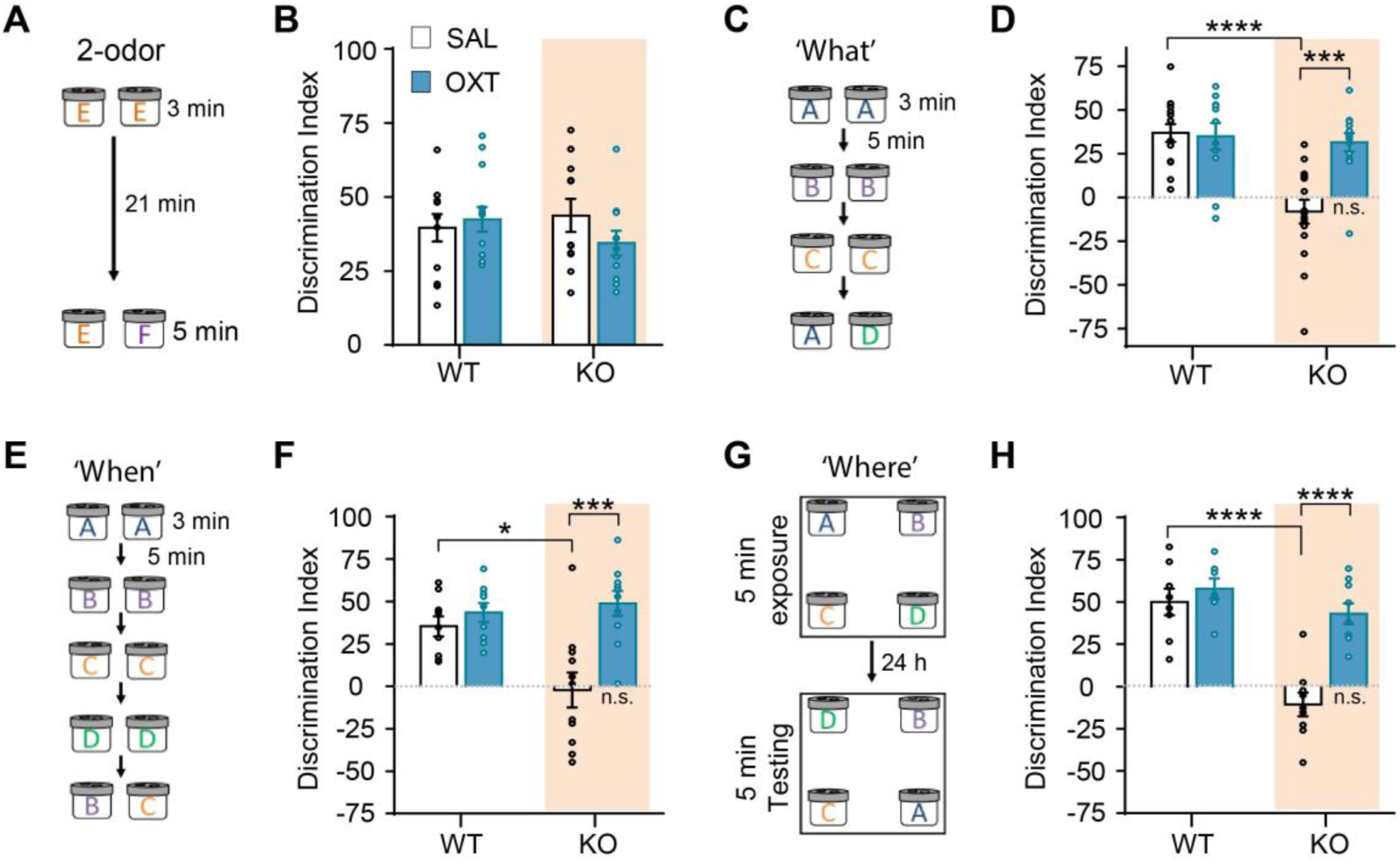
Episodic memory is restored in OXT-treated *Fmr1*-KO mice. Behavioral tests evaluated odor discrimination and acquisition of 3 components of episodic memory (‘what’, ‘where’, ‘when’) as described in Methods. (**A**) The 2-odor control task. (**B**) All groups preferentially discriminated novel odor F from the familiar odor E (F_(1,46)_=1.53, p=0.223). (**C**) Episodic ‘What’ task. (**D**) In the What task both WTs and *Fmr1*-KOs given postnatal OXT treatment preferentially explored novel odor D vs. familiar odor A, whereas KO-SAL mice did not (F_(1,50)_=10.2, p=0.002). (**E**) Episodic ‘When’ task. (**F**) *Fmr1*-KOs treated with SAL failed to discriminate earlier presented odor B vs. later presented odor C; in contrast, *Fmr1*-KOs given OXT preferentially explored less recently sampled odor B with discrimination indices comparable to the WT groups (F_(1,35)_=7.99, p=0.008). (**G**) Episodic ‘Where’ task. (**H**) Both WT groups preferentially explored the novel-location cues (A:D) without evident effect of OXT treatment. In contrast, SAL-treated KOs did not discriminate the switched odor pair whereas those given OXT preferentially explored the novel-location cues with discrimination indices comparable to the WT groups (F_(1,29)_=11.0, p=0.003). Statistics for all graphs: two-way ANOVA (interaction) with Tukey’s post-hoc comparisons indicated as: *p<0.05, ***p<0.001, ****p<0.0001, and n.s. p<0.05 for WT-OXT vs. KO-OXT. Legend in panel B also applies to panels D,F, and H.

Next, we evaluated encoding of the locations of cues and the order in which they were sampled (episodic ‘where’ and ‘when’, respectfully). To assess ‘when’, mice were exposed to a sequence of odor pairs (A-A, B-B, C-C, D-D), followed by a pair including odors B and C (**Fig. 2E**). WT mice (±OXT) preferentially explored the less recently encountered odor (B) whereas SAL-treated KOs did not; this deficit was fully corrected by OXT treatment (**Fig. 2F, S2C**). In the ‘where’ task, mice explored a chamber with 4 different odors presented simultaneously, and then 24-hrs later were tested in the same chamber with the positions of two diagonally opposed odors swapped (**Fig. 2G**). Both WT groups preferentially investigated the novel-location odors whereas SAL-treated KOs did not. However, *Fmr1*-KOs given OXT preferentially explored the relocated odors (**Fig. 2H, S2D**).

### Early-life OXT rescues hippocampal CA1 synaptic plasticity as assessed in adult Fmr1-KOs

Acquisition of the basic elements of episodic memory depends on hippocampus^25,26^, and *Fmr1*-KO mice exhibit significant LTP deficits in at least two of the links in the prime hippocampal circuit: the LPP innervation of the DG, and the CA3 to CA1, S-C projections^29,30^. It is possible that postnatal OXT rescues episodic memory in the KOs by restoring synaptic plasticity in these systems. We accordingly used hippocampal slices to evaluate the effects of early-life OXT treatments on basal synaptic transmission and LTP. Stimulation electrodes and recording electrodes were placed in CA1 str. radiatum to measure fEPSPs evoked by activation of the S-C afferents (**Fig. 3A**). Input/output curves were not detectably different for WT vs. *Fmr1*-KO slices (**Fig. 3B**), nor was the frequency facilitation elicited by a 10-pulse gamma frequency (40 Hz) (**Fig. 3C**). It thus appears that the mutation does not significantly alter basal transmission or the facilitated release produced by repetitive activation.

**Figure 3.**
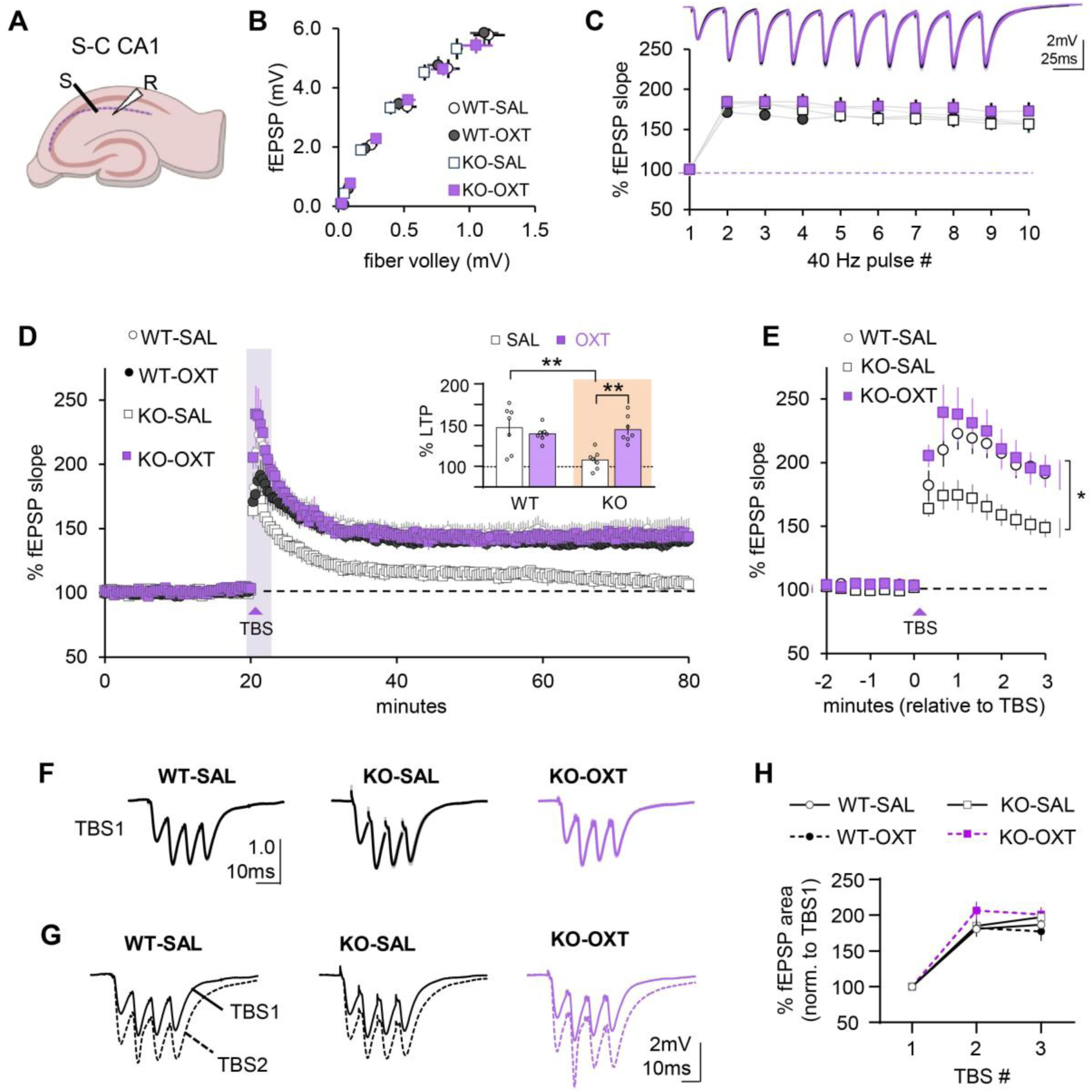
Postnatal Oxytocin treatment restores Schaffer-commissure (S-C), field CA1 LTP. (**A**) Acute hippocampal slices were prepared from adult male WT and *Fmr1*-KO mice treated daily with saline (SAL) or oxytocin (OXT) from P7 to P13. The S-C projections were stimulated (S) and field excitatory postsynaptic potentials (fEPSPs) were recorded (R) from CA1 Str. Radiatum; see Methods. (**B**) Input (fiber volley amplitude)/output (slope of the fEPSP) curves were similar across all groups, indicating no effects of genotype or treatment (F_(3,190)_=1.87, p=0.136; linear regression). (**C**) Responses to a ten pulse 40-Hz train were comparable across groups (F_(27,270)_=0.915, p=0.59); legend is same as in panel B. Superimposed traces (top) of mean (±SEM) fEPSP responses to the 40-Hz train for WT-SAL (black) and KO-SAL (purple) mice. (**D**) To assess S-C LTP, responses to 0.05 Hz stimulation were collected for 20-min before delivering a single train of 3 theta bursts (TBS). Both WT groups exhibited stable potentiation from 40-80 min post-TBS, whereas SAL-treated KOs declined towards baseline. In contrast, KO-OXT mice exhibited stable potentiation that was comparable to the WTs. *Inset bar graph:* Mean potentiation from 55-60 min post-TBS (F_(1,24)_=11.04, p=0.0028, 2-way ANOVA; **p<0.01 post-hoc Tukey). (**E**) Stretched axis from shaded area of panel D (min 18-23) shows that initial increases in fEPSP size after TBS are markedly reduced in KO-SAL mice but restored in KO-OXT mice, resembling WT-SAL profiles (F_(30,270)_=4.142, p<0.0001; post-hoc Tukey: *p<0.05 for WT-SAL and KO-OXT vs. KO-SAL). (**F**) Traces show mean (± SEM, n=7/group) responses to the first theta burst (TBS1), normalized to the amplitude of the response to first pulse. (**G**) Representative superimposed fEPSP responses to the first burst (TBS1, solid line) and to the subsequent burst (TBS2, dashed line), illustrating the response facilitation from TBS1 to TBS2. (**H**) Facilitation in fEPSP areas across the three bursts was comparable between groups (F_(6,48)_=1.467, p=0.2095); plots show (mean ± SEM) values. Statistics (C, E, and H): 2-way RM ANOVA (interaction) with post-hoc Tukey analyses.

Prior work showed that impairments to CA1 LTP in the *Fmr1*-KO are made evident by using near threshold levels of inducing stimulation^30^. We therefore used a minimum induction protocol involving a single train of three theta bursts (TBS; total of 12 stimulation pulses) which induced a sizeable degree of stable potentiation in WT slices but failed to elicit potentiation in KO slices. OXT administered from P7-P13 fully restored S-C LTP in adult *Fmr1*-KO slices, without effect on potentiation in WTs (**Fig. 3D**). This first use of the three-burst paradigm led to the unexpected observation that the magnitude of potentiation in WTs steadily increases across the three single stimulation pulses delivered in the minute following TBS. There was a near equivalent increase in the slope of the first pulse response after TBS in SAL-treated WTs and KOs (WT vs. KO: 181.9±11.9 vs. 163.7±6.3%; p=0.201, Tukey’s post-hoc) but the subsequent increases in each pulse response were significantly reduced in KOs relative to WTs (p<0.025, post-hoc). Early-life OXT treatment reinstated the post-induction facilitation effect in KO slices (**Fig. 3E**). In all, *Fmr1*-KO disrupts events occurring shortly after S-C LTP induction and this abnormality is stably corrected by OXT.

Next, we tested for KO-related perturbations to the theta-burst responses used to induce CA1 LTP. The digitized waveforms for each fEPSP in the initial TBS burst (4 pulse, 100 Hz) were expressed as a percent of the fEPSP amplitude of the first pulse after which a mean burst response was assembled for each slice (**Fig. 3F**) and then calculated average (±SEM) for all slices in three groups: WTs with early-life SAL (41.56±0.85); and *Fmr1*-KOs given early-life SAL (41.15±0.9) or OXT (39.64±2.18). The area of the fEPSP response to the first burst were comparable across all groups (One-way ANOVA (interaction): F_(2,19)_=0.5129, p=0.607). And, all three groups also showed a pronounced increase in the response areas between the first and subsequent bursts in a train (**Fig. 3G**). This effect is due in part to reduced shunting inhibition and the addition of NMDAR-gated potentials to the fEPSPs^50–52^. The three consecutive burst elicited comparable fEPSP areas across treatments and genotypes (**Fig 3H**).

Overall, these results show the *Fmr1*-KO condition does not measurably change synaptic responses evoked by theta bursts but nonetheless truncates the growth of potentiation in the post-induction period. This strongly suggests that the mutation leaves the initial and immediate induction steps intact but disrupts the normal development of LTP expression. Early-life OXT normalized the response growth after TBS and fully restored LTP in *Fmr1*-KOs without effect on WTs.

### Early life OXT restores LPP-LTP in adult Fmr1-KO

Evidence that LPP-LTP is impaired in *Fmr1*-KOs^29,53^ is of particular interest because the LPP-DG connection plays a vital role in episodic memory, including acquisition in the three episodic memory paradigms used here^38^. We tested if the P7-P13 OXT treatments that rescued episodic memory also normalize LPP-LTP (**Fig 4A**). LPP input/output curves were comparable in slices from WT and *Fmr1*-KO mice and, in both, were unaffected by OXT treatment (**Fig. 4B**). Gamma frequency stimulation of the LPP causes an initial within-train facilitation of DG fEPSPs followed by a steady decline in response size to below baseline values, an unusual response profile that reflects release variables^40^. This profile was intact in all four groups, suggesting that signal processing at the LPP-DG contact is not affected by genotype or postnatal OXT treatment (**Fig. 4C**). Despite baseline transmission being comparable across groups, LPP-LTP was reduced in SAL-treated KOs relative to SAL-treated WTs. Early-life OXT increased LPP-LTP magnitude in *Fmr1*-KOs to WT levels, without an effect on LTP in the WTs (**Fig. 4D,E**).

**Figure 4:**
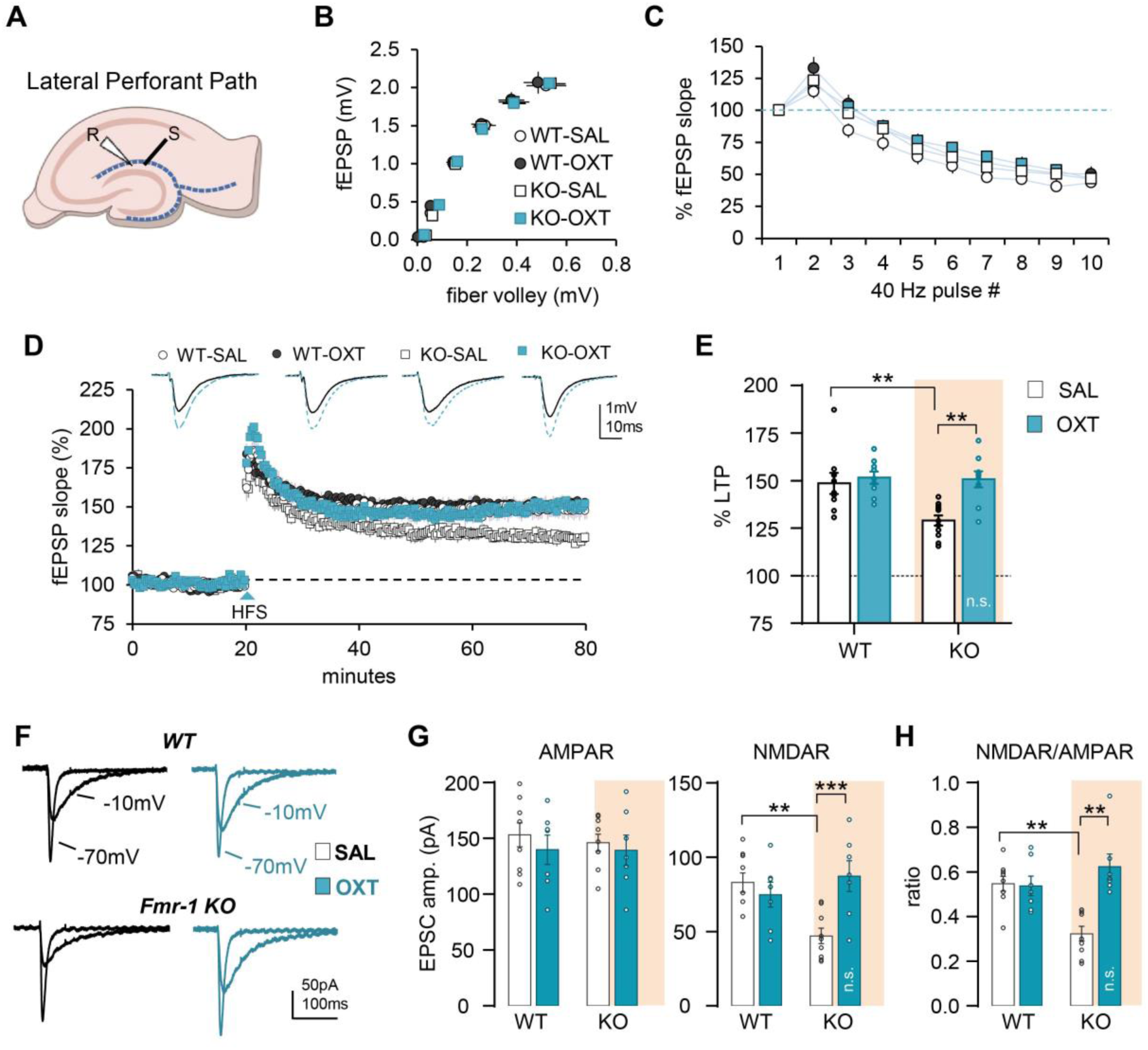
Early-life OXT treatment rescues lateral perforant path (LPP) LTP in *Fmr1*-KO mice. (**A**) Acute field recordings were collected from horizontal hippocampal slices of adult male WT and KO mice given daily SAL or OXT treatment from P7-P14; LPP (dashed blue line) stimulation (S) and recording (R) electrodes were placed in the dentate gyrus outer molecular layer. (**B**) Input/output curves were comparable across all groups (F_(3,118)_=0.109, p=0.955; linear regression). **C**) LPP stimulation with a ten-pulse 40-Hz train elicited a similar fEPSP response profile (F_(27,234)_=1.28, p=0.171; two-way RM ANOVA). (**D**) Following 20 min of baseline recording, LPP-LTP was induced by delivering a single 1-sec 100 Hz train (HFS). Stable potentiation was observed in both WT groups. *Fmr1*-KOs that received SAL showed impairment in LPP-LTP whereas KOs given postnatal OXT treatment exhibited LPP-LTP at WT levels. Representative traces show pre-HFS (black) and 79-80 min post-HFS fEPSP responses (blue) for each group. (**E**) Mean potentiation at 75-80 min post-HFS (from panel D; F_(1,34)_=5.61, p=0.024). (**F-G**) Granule cell whole cell recordings were used to assess NMDAR/AMPAR current ratios with LPP stimulation. (**F**) Representative EPSC traces with the membrane potential held at −10mV and −70mV for WT and KO groups. (**G**) AMPAR currents were similar across all groups (F_(1,28)_=0.086, p=0.77) In contrast, NMDAR-mediated currents were markedly lower in KO-SAL vs. WT-SAL, but restored by OXT treatment to WT levels (F_(1,28)_=10.3, p=0.003). (**H**) The AMPAR to NMDAR evoked current ratio was comparable in the two WT groups, markedly low in the KO-SAL group, and restored to WT levels in the KO-OXT group (F_(1,28)_=14.5, p=0.0007). Statistics (E, G, and H): 2-way ANOVA (interaction) with post-hoc Tukey comparisons (**p<0.01, ***p<0.001; n.s. for p >0.05 for WT-OXT vs. KO-OXT). Legend in panel E also applies to panels G and H.

Previous work determined that two factors were primarily responsible for the loss of LPP-LTP in the KOs: 1) a reduction in NMDAR-mediated synaptic responses, and 2) a loss of the endocannabinoid signaling needed to trigger the presynaptic changes that express potentiation at LPP-DG synapses^43,53,54^. Here we tested if early-life OXT treatment normalizes evoked NMDAR currents in the LPP-DG connection of *Fmr1*-KOs. Whole-cell recordings confirmed that AMPAR-gated LPP-EPSCs were comparable in slices from adult WT and *Fmr1*-KO mice given either OXT or SAL from P7-P13 (**Fig. 4F**, **4G**). In contrast, NMDAR-mediated currents were markedly reduced in SAL-treated KOs as compared to WT mice (**Fig 4G**). Early-life OXT restored NMDAR-mediated synaptic responses in KOs to WT levels, and consequently normalized the NMDAR/AMPAR current ratio in the mutants (**Fig 4H**). These results describe a specific modification produced by early-life OXT that is closely related to both physiological (LPP-LTP) and behavioral (episodic memory) outcomes.

### Effects of OXT given during adolescence

The hypothesis that OXT treatment normalizes an otherwise aberrant developmental trajectory associated with FXS gives rise to the prediction that the ameliorating effects of OXT will depend on the age at which it is administered. To test this, a separate cohort of mice was given intranasal OXT or SAL during adolescence (daily from P30-P36), and assayed as adults for LTP and episodic memory (**Fig. 5A**). Mice in all four groups (WT±OXT; KO±OXT) had high retention scores in the two-odor discrimination test with no measurable effects of genotype or treatment (**Fig. 5B, S3A**). The SAL-treated KOs again had a severe deficit in the episodic ‘what’ task and, unlike the case for early-life administration, this was not restored by adolescent OXT treatment; there was no treatment effect on WT performance (**Fig. 5C, S3B**). Adolescent OXT also failed to offset the LPP-LTP impairment in adult *Fmr1*-KOs (**Fig. 5D,E**). In all, the delayed P30-P36 OXT treatment had no evident effects on episodic memory or the plasticity needed for its encoding.

**Figure 5:**
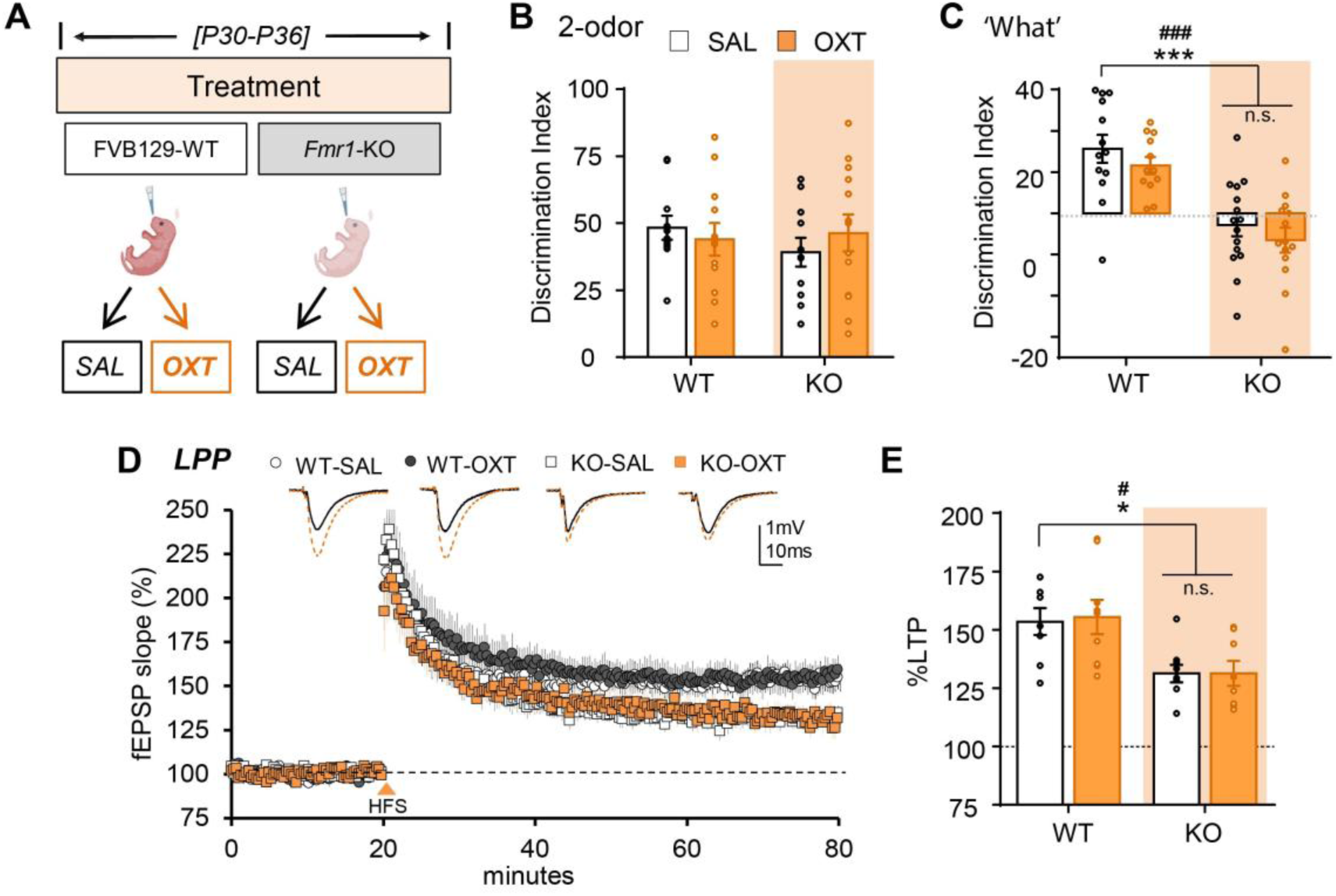
OXT treatment during adolescence does not restore learning or LTP in Fmr1-KOs. (**A**) WT and *Fmr1*-KO mice received daily intranasal OXT or SAL treatment from P30 to P36. Effects of treatment were assessed at 2-4 months of age. (**B**) In the 2-odor discrimination task, all groups successfully discriminated the novel odor after a delay (see Fig. 2A) (Interaction: F_(1,45)_=0.643, p=0.427). (**C**) In the episodic ‘What’ task (see Fig. 2C), WT mice given SAL or OXT preferentially explored the novel odor. In contrast, *Fmr1*-KOs given SAL or OXT failed to show a preference (Interaction: F_(1,50)_=0.003, p=0.959; Genotype: F_(1,50)_=40.0, p<0.0001; post-hoc Tukey: ***p=0.0002 KO-SAL vs. WT-SAL, ###p=0.0004 KO-OXT vs. WT-SAL, n.s. p=0.78 for KO-SAL vs. KO-OXT) (**D, E**) The effect of P30-P36 OXT treatment on LPP-LTP was evaluated in adult hippocampal slices. (**D**) Stable and comparable LPP-LTP was observed in slices from WT mice given SAL or OXT during adolescence whereas potentiation was significantly reduced in KOs independent of treatment. Representative traces show pre-HFS (black) and 60-min post-HFS (orange) responses for reach group. (**E**) Mean LPP potentiation at 55-60 min post-HFS (from panel D; Interaction: F_(1,30)_=0.028, p=0.868; Genotype: F_(1,30)_=16.55, p=0.0003; post-hoc Tukey: *p=0.046 KO-SAL vs. WT-SAL, #p=0.0265 KO-OXT vs. WT-OXT, n.s. p>0.999 KO-SAL vs. KO-OXT). Statistics (B, C, and E): 2-way ANOVA with Tukey’s post-hoc analyses. Legend in panel B also applies to panels C and E.

### Effects of acute OXT treatment on defects in two forms of LTP

Evidence that OXT expression is abnormally low in *Fmr1-*KOs^17,18^ and that postnatal OXT treatment increases OXT expression in the PVN^19^, raises the possibility that treatment-related increases in endogenous OXT efflux normalize plasticity in adult *Fmr1*-KOs. This argument predicts that acute OXT infusion will reproduce the effects of early life OXT treatments on plasticity in the adult *Fmr1*-KO hippocampus. Tests of this possibility produced an intriguing set of results. Acute OXT treatment did not change the magnitude of LPP-LTP in *Fmr1*-KO slices relative to vehicle-treated KO slices (**Fig. 6A**). OXT infusion also did not influence LPP-LTP in WTs. As described, we confirmed earlier reports^29,53,55,56^ that NMDAR-gated EPSCs are markedly reduced in *Fmr1*-KO LPP-DG synapses. This disturbance along with the defect in LPP-LTP was corrected by early-life OXT (see Fig 4G). However, acute OXT infusion into adult KO slices had no measureable effect on the magnitude of NMDAR-gated evoked responses (**Fig. 6B,C**) or on the NMDAR/AMPAR response ratio **(Fig. 6D**). Thus, for LPP-DG synapses, acute OXT treatment did not produce the recovery of the normal LTP and NMDAR operations that were found with early-life OXT administration.

**Figure 6:**
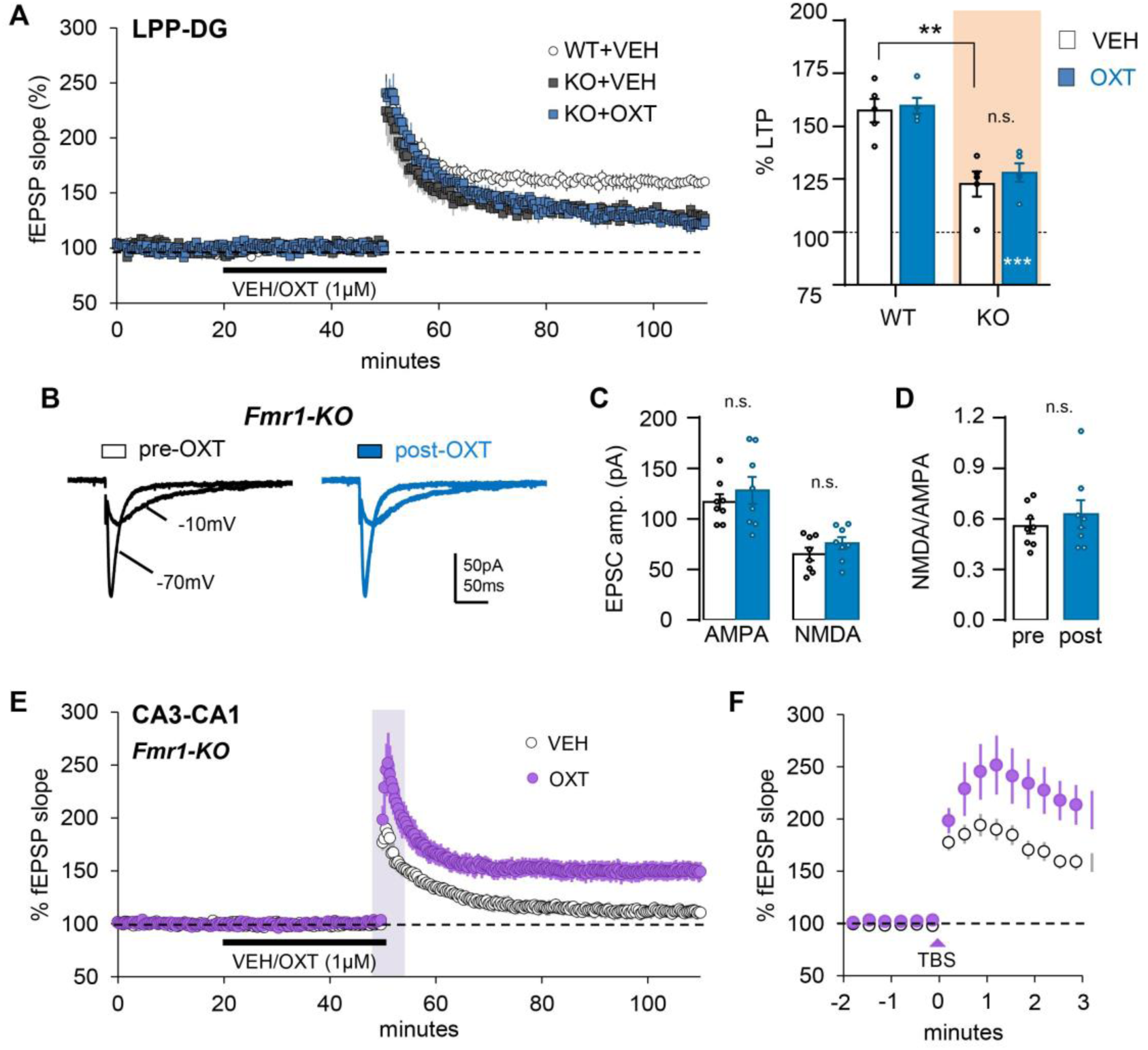
Acute OXT infusion increases S-C LTP in *Fmr1*-KOs but does not normalize measures of LPP function. (**A**) Horizontal hippocampal slices from naive WT and KO mice were infused with vehicle (VEH) or OXT (1-µM) for 30 min before being given HFS to induce LPP-LTP. *Left*: *Fmr1*-KO slices treated with VEH have impaired LPP-LTP relative to WT+VEH slices and potentiation in the KOs is not influenced by OXT infusion. *Right.* Bar graph summarizing the percent LTP at 105-110 min shows that KO+VEH slices exhibit a significant impairment in the magnitude of LPP-LTP as compared to WT-VEH measures (Interaction: F_(1,16)_=0.093, p=0.764; Genotype: F_(1,16)_=45.08, p<0.0001; post-hoc Tukey: **p=0.0017 KO+VEH vs. WT+OXT); infusion of OXT did not facilitate LPP-LTP in KO slicees (n.s. p=0.870 KO+VEH vs. KO+OXT). (**B**) Whole-cell recordings of the granule cells: EPSCs were elicited by LPP stimulation in slices from *Fmr1*-KOs and were held at −10mV to measure NMDAR currents or −70mV for AMPAR currents. (**C**) There was no effect of OXT treatment on LPP AMPAR EPSCs (p=0.2254, 2-tailed paired t-test), NMDAR-currents (p=0.0546) or (**D**) the NMDAR/AMPAR-mediated current ratio (p=0.3547). (**E,F**) CA1 str. radiatum field recordings of S-C responses in *Fmr1*-KO slices show that acute OXT infusion increased the magnitude of S-C LTP (**E**) and robustly enhanced the increase in fEPSP amplitude across the first 3 min post-TBS (**F**, panel shows an expansion of the X-axis from 48-53 in panel E; Interaction: F_(16,192)_ = 5.916, p<0.0001, 2-way RM ANOVA).

In marked contrast to the LPP results, acute OXT infusion fully normalized CA1-LTP elicited by three theta bursts in adult *Fmr1*-KO slices (**Fig. 6E**). The early post-TBS growth of fEPSPs otherwise impaired in the *Fmr1*-KOs (see Fig. 3E) was also corrected by acute OXT infusion (**Fig. 6F**).

## DISCUSSION

Several studies suggest that administration of OXT early in postnatal life can correct sociability deficits in mouse models of autism^17,19^. This result is of considerable importance because disruption of social interactions is a prominent and debilitating problem for autistic individuals^57^. The present studies show that the positive effects of early-life OXT treatment extend to cognitive difficulties that are a salient feature of autism and other neurodevelopmental disorders^58–60^. Specifically, we show that spatial learning and the acquisition of three basic elements of episodic memory -- a form of encoding that is vital for orderly thinking^20,60–62^ are severely impaired in *Fmr1*-KOs but fully restored by early-life OXT treatment. Defects in two forms of memory-related synaptic plasticity were also corrected by the early-life OXT treatments. Given that the hippocampus plays a central role in episodic memory^21,26,61,63^, it is likely that OXT’s effects on the synaptic defects are major contributors to the behavioral recovery.

While rodents presumably lack the narrative and autobiographical components of human episodic memory, there is increasing evidence that they utilize a rudimentary but analogous type of encoding^61,64,65^ which is critically dependent on the hippocampus^38^. Specifically, rats and mice retain information about the identities, locations, and sequence (episodic ‘what’, ‘where’, and ‘when’) for items encountered during exploration of a novel environment^64,66–68^. Learning thus occurs without rewards or practice and has a critical temporal dimension -- these are essential components of human episodic memories. Failures in this form of encoding are thought to be important contributors to the intellectual disabilities that characterize FXS and ASD and, importantly, we found that *Fmr1*-KOs exhibit profound deficits in all three elements of episodic memory. Like humans, rodent episodes are also strongly associated with the context in which they were acquired^69^. Whether this aspect of encoding is also affected in the mutants remains to be determined.

That early-life OXT normalized LTP in *both* CA1 and LPP of *Fmr1*-KOs was surprising because of the substantial differences in the cellular substrates for the two forms of plasticity. The loss of stable potentiation in mutant LPP-DG synapses is due to impairments to NMDAR-gated synaptic currents and retrograde (spine to terminal) endocannabinoid signaling^43^. We confirmed here that LPP-evoked NMDAR-, but not AMPAR-, mediated responses are reduced in *Fmr1*-KOs and that early-life OXT treatment produces a remarkably complete normalization of these synaptic currents. How OXT elicits this effect is unclear. A reduction in NMDAR currents was not detected for a second afferent of the granule cell dendrites (i.e., the medial perforant path) in earlier work^29^, suggesting that the defect in the KOs reflects disruption of orthograde (terminal to spine) signaling. NMDARs are targeted by multiple kinases and associate with a large number of signaling proteins^70^. Perturbations to these activity-dependent relationships could alter NMDAR operations in the afferent specific manner found in *Fmr1*-KOs. As such, rescue by postnatal OXT treatment might reflect an influence on the development of a particular relationship between LPP terminals and postsynaptic NMDARs.

An argument of the above kind is not applicable to the early-life OXT driven recovery of CA1-LTP. Earlier work found no evidence for a loss of NMDAR signaling at CA3-CA1 contacts in *Fmr1*-KOs^53^. Rescue of NMDAR-dependent synaptic plasticity in *Fmr1*-KO mice^30^ and the theta burst responses recorded in the present experiments appeared to be of normal (WT) size in CA1. There was, however, an evident difference between genotypes for synaptic responses during the first minute post-TBS. The use of threshold induction conditions uncovered what appears to be a previously unrecognized growth phase in fEPSPs during this period in WTs and a truncation of this effect in the mutants. It is generally agreed that CA1-LTP is expressed by an increase in postsynaptic AMPARs^71^ and AMPAR currents^72–74^. Relatedly, potentiation is associated with expansion of the activated spine ^75,76^ and its postsynaptic density^77^. Facilitation at the 20 sec point following TBS is all but completely eliminated by NMDAR antagonists^69^ indicating that response enhancement is due to postsynaptic events even at this early time point. Collectively, these observations point to a hypothesis in which the mutation disrupts the development of processes that mediate TBS-driven structural modifications to the postsynaptic compartment and/or migration of AMPARs into the newly enlarged receptor zone. In any event, early-life OXT restored the progressive increase of synaptic responses, suggesting that OXT corrects developmental problems relating to the initial shift of synapses into their potentiated state.

Acute OXT infusion revealed distinct effects on the two types of LTP impairments observed in the *Fmr1-*KO model. Infusion produced a complete recovery in CA1 including restoration of the growth of potentiation immediately following TBS. However, in the LPP pathway, acute OXT infusion did not significantly affect LTP or the associated NMDAR hypofunction. Studies have shown that hypothalamic OXT expression is reduced in both *Fmr1* and *Cntnap*2 knockout mice, and that levels of the hormone are normalized by early postnatal OXT treatment^17,19^. While the effect of acute OXT on CA1-LTP suggests that postnatal treatment could potentially restore OXT supply to the KO hippocampus and preserve LTP capability, this seems unlikely. Any native OXT in hippocampus would presumably be removed by the two hours of washout that preceded physiological testing. But it is possible that the cell biological effects of the hormone persist and thus are still present in slices at the start of LTP experiments. This argument implies a rapid onset for the LTP-related cellular process targeted by OXT to account for the recovery produced by infusion into adult KO slices.

The lack of LPP-LTP rescue with acute OXT infusion does not necessarily rule out the possibility that the positive effects of early-life treatments were mediated by OXT up-regulation. OXT’s influence on NMDAR operations in LPP-DG synapses could be slow to develop and thus may not be evident following the brief applications evaluated here. It will be interesting in future studies to determine if prolonged infusions have greater effect on LPP-LTP.

An alternative to the ‘restoration of OXT signaling’ hypothesis is that the corrective effects of early-life OXT treatment on LTP are due to normalization of developmental trajectories that are essential for synaptic plasticity and associated memory functions. This would be consistent with our finding that later adolescent OXT treatment had no effect on plasticity or learning in the KOs. Moreover, OXT receptors are expressed at high levels in neocortex and hippocampus from P7 to P21^33,78^, a period that overlaps the present treatment timeline. Thus, early-life OXT treatment may compensate for low levels of the endogenous neuropeptide in the KOs to facilitate the emergence of LTP early in the second postnatal week^79,80^. OXT has potent effects on hippocampal dendrites and spines during this period whereas genetic deletion of the OXT receptor causes marked changes to spine morphology and number^81^. Possibly, then, the developmental steps that generate the pre- and post-synaptic interactions underlying LPP-LTP require OXT signaling and, having failed to emerge in the KO, cannot be substituted for by either adolescent or adult applications. It should be noted that such developmental trajectory arguments could apply equally well to the recovery of LTP in CA3-CA1 synapses.

Further development of the above ideas requires additional information on the defects contributing to impaired LTP in *Fmr1*-KOs and the downstream signaling of the OXT receptor. Our prior work on CA1-LTP showed that the complex machinery responsible for activity driven actin polymerization, an event required for LTP consolidation, is not impaired in the mutants. However, signaling cascades involving the small Rho GTPase Rac and other proteins that stabilize newly formed actin filaments are defective in the *Fmr1*-KOs^82,83^. OXT is known to engage Rho-GTPases in various cell types, and thereby promotes actin network reorganization^84,85^. Whether OXT regulates the actin cytoskeleton at CA3-CA1 synapses, and early-life treatment restores actin signaling cascades in *Fmr1*-KO mice, remains to be resolved. It is difficult to pose specific hypotheses about the manner in which early-life OXT restores NMDAR functions at the LPP-DG synapses without first explaining how the mutation selectively disrupts receptor operations at this site. Clarification of these issues could lead to broader hypotheses about OXT’s roles in the normal development and later operation of cortical networks.

Overall, the present findings build on prior work on early-life OXT treatment to show that in *Fmr1*-KO mice, treatments limited to the second postnatal week have enduring normalizing effects on both social behavior and the acquisition of hippocampus-dependent spatial and episodic memories. These behavioral effects were associated with a restoration of distinct forms of LTP within two populations of synapses in hippocampus^26^. These results support the general idea that early life intervention during childhood may be critical for treating cognitive dysfunction in FXS and other autism associated disorders.

## ACKNOWLEDGEMENTS

The authors thank Ms. Lida Aghazadah for contributions to behavioral analyses and Ms. Lucy Yao for technical assistance.

## AUTHOR CONTRIBUTIONS

BMC, JCL, GL, and CMG conceived and designed the studies; GL and CMG supervised the project. JC, AAL, and BMC performed and analyzed the behavioral experiments; JC, AAL and YJ performed and analyzed the electrophysiological experiments; JC, AAL, JCL, GL, and CMG wrote the manuscript. All of the authors read and accepted the manuscript.

