## Supplementary Materials for "Early-life Oxytocin Rescues Hippocampal Synaptic Plasticity and Episodic Memory in a Mouse Model of Fragile X Syndrome"

### Supplementary Material

Table S1. List of odorants used in behavioral tasks

| Odorant ID | Odorant (name, company) | Concentration (odorant: mineral oil) |
|---|---|---|
| A | (+) -Limonene (>97% purity, *Sigma- Aldrich*) | 1:4000 |
| B | Cyclohexyl ethyl acetate (>97%, *International Flavors & Fragrances Inc.*) | 1.97:4000 |
| C | Citronellal 96% (~96%, *Alfa Aesar*) | 1.5:4000 |
| D | Octyl aldehyde 99% (~99% *Acros Organics*) | 1.5:4000 |
| E | Anisole 99% (~99% *Acros Organics*) | 0.85:4000 |
| F | 1-Pentanol 99% (~99% *Acros Organics*) | 1.36:4000 |

*Each odorant was dissolved in mineral oil (~0.1 Pa).

Table S2. Statistical values for behavioral analyses.

**Treatment P7-P13**

| Fig | | | Mean±SEM | Two-way ANOVA |
|---|---|---|---|---|
| **1C** | Social Approach | Object vs S1 | WT-SAL: 49.2±5.8<br>WT-OXT: 56.1±4.3<br>KO-SAL: 55.3±5.9<br>KO-OXT: 67.7±4.9 | Interaction: $F_{(1,40)}$=0.25, p=0.62<br>Genotype: $F_{(1,40)}$=2.68, p=0.11<br>Treatment: $F_{(1,40)}$=3.16, p=0.08 |
| | | | | Post hoc Tukey |
| | | | | WT-SAL vs. WT-OXT: p=0.78<br>WT-SAL vs. KO-SAL: p=0.83<br>WT-SAL vs. KO-OXT: p=0.09<br>WT-OXT vs. KO-SAL: p=0.99<br>WT-OXT vs. KO-OXT: p=0.48<br>KO-SAL vs. KO-OXT: p=0.42 |
| **1D** | Social Recognition | S1 vs S2 | WT-SAL: 28.9±7.8<br>WT-OXT: 25.1±6.8<br>KO-SAL: -0.97±7.8<br>KO-OXT: 28.1±4.1 | Interaction: $F_{(1,37)}$=5.15, p=0.03<br>Genotype: $F_{(1,37)}$=3.46, p=0.07<br>Treatment: $F_{(1,37)}$=3.0, p=0.09 |
| | | | | Post hoc Tukey |
| | | | | WT-SAL vs. WT-OXT: p=0.98<br>WT-SAL vs. KO-SAL: *p=0.02<br>WT-SAL vs. KO-OXT: p=0.99<br>WT-OXT vs. KO-SAL: p=0.054<br>WT-OXT vs. KO-OXT: p=0.99<br>KO-SAL vs. KO-OXT: *p=0.047 |
| **1F** | OLM | Stationary vs moved | WT-SAL: 25.4±5.1<br>WT-OXT: 20.2±6.6<br>KO-SAL: -2.2±2.4<br>KO-OXT: 26.6±4.3 | Interaction: $F_{(1,40)}$=11.63, p=0.002<br>Genotype: $F_{(1,40)}$=4.48, p=0.04<br>Treatment: $F_{(1,40)}$=5.57, p=0.02 |
| | | | | Post hoc Tukey |
| | | | | WT-SAL vs. WT-OXT: p=0.87<br>WT-SAL vs. KO-SAL:** p=0.002<br>WT-SAL vs. KO-OXT: p=0.99<br>WT-OXT vs. KO-SAL: *p=0.012<br>WT-OXT vs. KO-OXT: p=0.80<br>KO-SAL vs. KO-OXT: **p=0.0015 |
| **1G** | Distance Travelled | Empty | WT-SAL:17.1±0.80<br>WT-OXT:15.0±0.63<br>KO-SAL: 20.4±0.9<br>KO-OXT: 18.4±0.68 | Interaction: $F_{(1,42)}$=0.0081, p=0.93<br>Genotype: $F_{(1,42)}$=18.8, p<0.0001<br>Treatment: $F_{(1,42)}$=7.17, p=0.011 |
| | | | | Post hoc Tukey |
| | | | | WT-SAL vs. WT-OXT: p=0.20<br>WT-SAL vs. KO-SAL: *p=0.19<br>WT-SAL vs. KO-OXT: p=0.66<br>WT-OXT vs. KO-SAL: ****p<0.0001<br>WT-OXT vs. KO-OXT: *p=0.020<br>KO-SAL vs. KO-OXT: p=0.29 |
| | | Exposure | WT-SAL: 22.1±1.9<br>WT-OXT: 19.2±1.6<br>KO-SAL: 24.8±2.4<br>KO-OXT: 24.5±1.1 | Interaction: $F_{(1,42)}$=0.52, p=0.48<br>Genotype: $F_{(1,42)}$=4.64, p=0.04<br>Treatment: $F_{(1,42)}$=0.76, p=0.39 |
| | | | | Post hoc Tukey |
| | | | | WT-SAL vs. WT-OXT: p=0.66<br>WT-SAL vs. KO-SAL: p=0.73<br>WT-SAL vs. KO-OXT: p=0.81<br>WT-OXT vs. KO-SAL: p=0.14 |

| | | | | |
|---|---|---|---|---|
| | | | | WT-OXT vs. KO-OXT: p=0.21<br>KO-SAL vs. KO-OXT: p=0.99 |
| | | Test | WT-SAL: 17.2±1.6<br>WT-OXT: 15.5±1.3<br>KO-SAL: 23.2±1.7<br>KO-OXT: 22.4±1.4 | Interaction: $F_{(1,42)}$=0.08, p=0.77<br>Genotype: $F_{(1,42)}$=17.5, p=0.0001<br>Treatment: $F_{(1,42)}$=0.64, p=0.43 |
| | | | | Post hoc Tukey |
| | | | | WT-SAL vs. WT-OXT: p=0.86<br>WT-SAL vs. KO-SAL: *p=0.035<br>WT-SAL vs. KO-OXT: p=0.11<br>WT-OXT vs. KO-SAL: **p=0.0043<br>WT-OXT vs. KO-OXT: *p=0.018<br>KO-SAL vs. KO-OXT: p= |
| **2B** | 2-odor | Familiar vs novel | WT-SAL: 39.6±4.6<br>WT-OXT: 42.5±4.2<br>KO-SAL: 43.7±5.6<br>KO-OXT: 34.4±4.2 | Interaction: $F_{(1,43)}$=1.72, p=0.20<br>Genotype: $F_{(1,43)}$=0.18, p=0.68<br>Treatment: $F_{(1,43)}$=0.48, p=0.49 |
| | | | | Post hoc Tukey |
| | | | | WT-SAL vs. WT-OXT: p=0.97<br>WT-SAL vs. KO-SAL: p=0.92<br>WT-SAL vs. KO-OXT: p=0.86<br>WT-OXT vs. KO-SAL: p=0.99<br>WT-OXT vs. KO-OXT: p=0.61<br>KO-SAL vs. KO-OXT: p=0.52 |
| **2D** | What | Familiar vs novel | WT-SAL: 37.0±5.1<br>WT-OXT: 35.1±7.6<br>KO-SAL: -8.0±6.8<br>KO-OXT: 31.7±5.3 | Interaction: $F_{(1,50)}$=10.8, p=0.002<br>Genotype: $F_{(1,50)}$=14.6, p=0.0004<br>Treatment: $F_{(1,50)}$=8.91, p=0.004 |
| | | | | Post hoc Tukey |
| | | | | WT-SAL vs. WT-OXT: p=0.99<br>WT-SAL vs. KO-SAL:****p<0.0001<br>WT-SAL vs. KO-OXT: p=0.93<br>WT-OXT vs. KO-SAL:****p<0.0001<br>WT-OXT vs. KO-OXT: p=0.98<br>KO-SAL vs. KO-OXT: ***p=0.002 |
| **2F** | When | Recent vs older<br>*(odor B vs odorC) | WT-SAL: 35.4±5.9<br>WT-OXT: 43.4±5.6<br>KO-SAL: -2.2±10.3<br>KO-OXT: 48.9±7.5 | Interaction: $F_{(1,35)}$=7.34, p=0.01<br>Genotype: $F_{(1,35)}$=4.1, p=0.05<br>Treatment: $F_{(1,35)}$=13.9, p=0.0007 |
| | | | | Post hoc Tukey |
| | | | | WT-SAL vs. WT-OXT: p=0.90<br>WT-SAL vs. KO-SAL: **p=0.0092<br>WT-SAL vs. KO-OXT: p=0.64<br>WT-OXT vs. KO-SAL: **p=0.0013<br>WT-OXT vs. KO-OXT: p=0.96<br>KO-SAL vs. KO-OXT: ***p=0.0002 |
| **2H** | Where | Stationary vs switched | WT-SAL: 50.0±7.9<br>WT-OXT: 57.8±6.1<br>KO-SAL: -10.6±7.1<br>KO-OXT: 43.0±6.1 | Interaction: $F_{(1,29)}$=11.0, p=0.003<br>Genotype: $F_{(1,29)}$=29.8, p<0.0001<br>Treatment: $F_{(1,29)}$=19.8, p=0.0001 |
| | | | | Post hoc Tukey |
| | | | | WT-SAL vs. WT-OXT: p=0.87<br>WT-SAL vs. KO-SAL: ****p<0.0001<br>WT-SAL vs. KO-OXT: p=0.88<br>WT-OXT vs. KO-SAL: ****p<0.0001<br>WT-OXT vs. KO-OXT: p=0.46<br>KO-SAL vs. KO-OXT: ****p<0.0001 |
| | | **Treatment P30-P36** | | |
| **5B** | 2-odor | Familiar vs novel | WT-SAL: 48.3±4.5<br>WT-OXT: 44.0±6.1<br>KO-SAL: 39.2±5.4 | Interaction: $F_{(1,43)}$=0.93, p=0.34<br>Genotype: $F_{(1,43)}$=0.32, p=0.58<br>Treatment: $F_{(1,43)}$=0.06, p=0.82 |

| | | | | |
|---|---|---|---|---|
| | | | KO-OXT: 46.4±6.9 | |
| | | | | Post hoc Tukey |
| | | | | WT-SAL vs. WT-OXT: p=0.96<br>WT-SAL vs. KO-SAL: p=0.72<br>WT-SAL vs. KO-OXT: p=0.99<br>WT-OXT vs. KO-SAL: p=0.94<br>WT-OXT vs. KO-OXT: p=0.99<br>KO-SAL vs. KO-OXT: p=0.83 |
| **5C** | What | Familiar vs novel | WT-SAL: 31.2±6.8<br>WT-OXT: 23.2±4.1<br>KO-SAL: -5.6±5.6<br>KO-OXT: -13.0±6.0 | Interaction: $F_{(1,50)}$=0.003, p=0.96<br>Genotype: $F_{(1,50)}$=40.0, p<0.0001<br>Treatment: $F_{(1,50)}$=1.80, p=0.19 |
| | | | | Post hoc Tukey |
| | | | | WT-SAL vs. WT-OXT: p=0.78<br>WT-SAL vs. KO-SAL: ***p=0.0002<br>WT-SAL vs. KO-OXT: ****p<0.0001<br>WT-OXT vs. KO-SAL: **p=0.0051<br>WT-OXT vs. KO-OXT: ***p=0.0004<br>KO-SAL vs. KO-OXT: p=0.78 |

*Red: preferred object or odor .

Table S3. Statistical values for raw supplementary data behavioral analyses.

| Fig | | | | Paired t-test |
|---|---|---|---|---|
| | | | | **Treatment P7-P13** |
| **S1A** | Social Approach | Object vs S1 | WT-SAL | $t_{11}$=6.606, p<0.0001 |
| | | | WT-OXT | $t_{10}$=7.724, p<0.0001 |
| | | | KO-SAL | $t_{10}$=4.193, p=0.0018 |
| | | | KO-OXT | $t_{8}$=9.692, p<0.0001 |
| **S1B** | Social Recognition | S1 vs S2 | WT-SAL | $t_{11}$=3.225, p=0.0079 |
| | | | WT-OXT | $t_{10}$=3.355, p=0.0073 |
| | | | KO-SAL | $t_{10}$=0.6457, p=0.5330 |
| | | | KO-OXT | $t_{7}$=2.711, p=0.0302 |
| **S1C** | OLM | Stationary vs moved | WT-SAL | $t_{10}$=4.854, p=0.0007 |
| | | | WT-OXT | $t_{11}$=2.895, p=0.0146 |
| | | | KO-SAL | $t_{10}$=0.8717, p=0.4038 |
| | | | KO-OXT | $t_{9}$=2.417, p=0.0388 |
| **S2A** | 2-odor | Familiar vs novel | WT-SAL | $t_{11}$=6.287, p<0.0001 |
| | | | WT-OXT | $t_{12}$=6.368, p<0.0001 |
| | | | KO-SAL | $t_{10}$=5.736, p=0.0002 |
| | | | KO-OXT | $t_{10}$=6.128, p=0.0001 |
| **S2B** | What | Familiar vs novel | WT-SAL | $t_{13}$=4.412, p=0.0007 |
| | | | WT-OXT | $t_{10}$=3.027, p=0.0127 |
| | | | KO-SAL | $t_{15}$=1.801, p=0.0919 |
| | | | KO-OXT | $t_{12}$=3.901, p=0.0021 |
| **S2C** | When | Recent vs older *(odor B vs odorC) | WT-SAL | $t_{8}$=2.762, p=0.0246 |
| | | | WT-OXT | $t_{8}$=5.468, p=0.0006 |
| | | | KO-SAL | $t_{11}$=0.3355, p=0.7676 |
| | | | KO-OXT | $t_{8}$=3.654, p=0.0065 |
| **S2D** | Where | Stationary vs switched | WT-SAL | $t_{7}$=3.221, p=0.0146 |
| | | | WT-OXT | $t_{6}$=6.706, p=0.0005 |
| | | | KO-SAL | $t_{8}$=0.8705, p=0.4094 |
| | | | KO-OXT | $t_{8}$=4.924, p=0.0012 |
| | | **Treatment P30-P36** | | |
| **S3A** | 2-odor | Familiar vs novel | WT-SAL | $t_{10}$=8.792, p<0.0001 |
| | | | WT-OXT | $t_{11}$=5.921, p=0.0001 |
| | | | KO-SAL | $t_{10}$=5.863, p=0.0002 |
| | | | KO-OXT | $t_{12}$=4.543, p=0.0007 |
| **S3B** | What | Familiar vs novel | WT-SAL | $t_{12}$=3.121, p=0.0088 |
| | | | WT-OXT | $t_{11}$=4.701, p=0.0006 |
| | | | KO-SAL | $t_{15}$=0.4782, p=0.6399 |
| | | | KO-OXT | $t_{13}$=1.208, p=0.2487 |

*Red: preferred object or odor . Within group comparisons: paired t-test (two-tailed)

### Figure S1

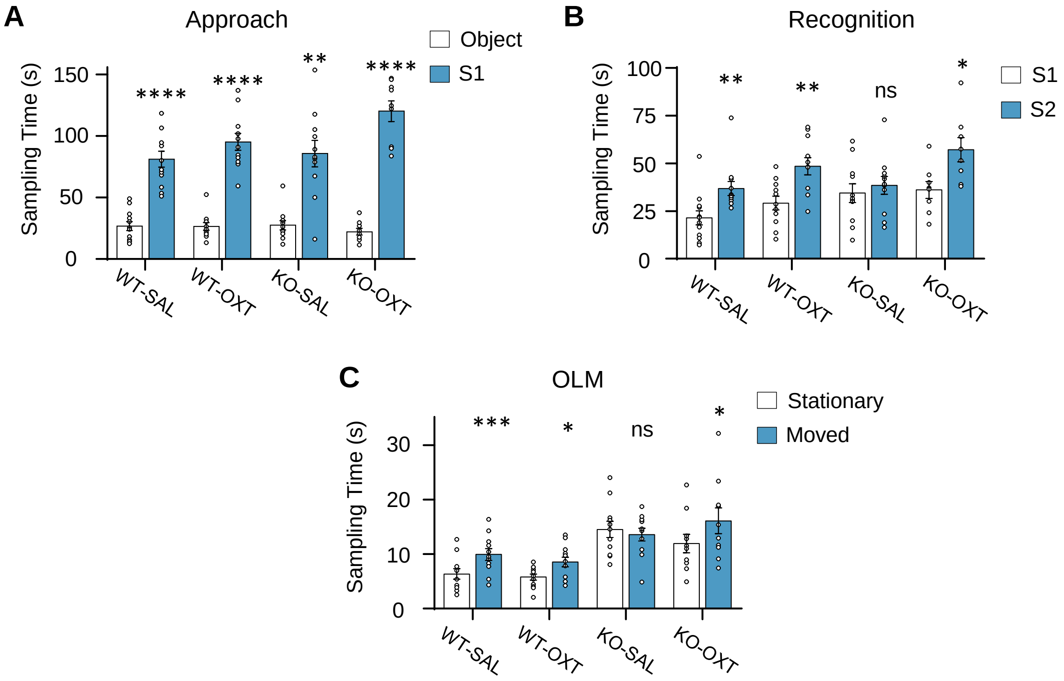


**Figure S1: Sampling times for three-chamber sociability task and object location memory (OLM) paradigms.** (**A, B**) Graphs show mean raw sampling time (seconds) used to calculate the discrimination indices for the three-chamber task during the Approach (see Fig. 1C) and Recognition (see Fig. 1D) phases. (**A**) *Approach:* Mean sampling time of the stranger mouse (S1) is greater than that of the object for all groups. (**B**) *Recognition*: Both WT groups (WT-SAL and WT-OXT) and KOs given OXT (KO-OXT) exhibited greater sampling times for the novel stranger mouse (S2) vs. the familiar mouse (S1). However, the KO-SAL group did not show a bias. (**C**) Graphs shows the mean raw sampling time for the OLM testing trial (see Fig. 1F). Time spent sampling the moved object vs. the stationary object was greater in both WT groups and OXT-treated KOs, whereas KO-SAL mice exhibited no preference for the moved object. *Statistics*: 2-tailed paired t-test: n.s. p>0.05, *p<0.05, **p<0.01, ***p<0.001, ****p<0.0001. Detailed statistical analyses are provided in supplementary table 3. Data presented as mean ± SEM.

### Figure S2

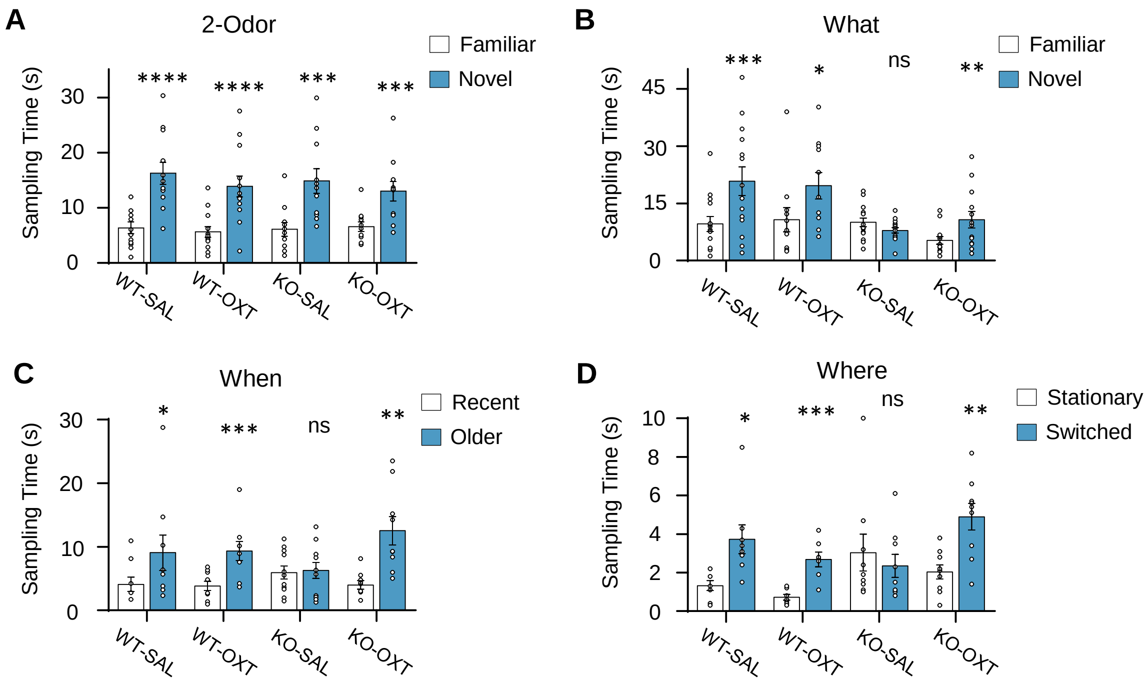


**Figure S2: Sampling times for 2-odor discrimination and episodic memory tasks.** (**A-D**) Graphs show the sampling times (seconds) of test odors used to calculate each animal's discrimination index presented in Fig. 2B (2-Odor), Fig. 2D ('What'), Fig. 2F ('When'), and Fig. 2H (Where). (**A**) *2-Odor:* All groups exhibited greater sampling times for the novel odor vs. the familiar odor. (**B**) *'What' task*: KOs given SAL (KO-SAL) had similar sampling times for the novel and familiar odors, whereas the other three groups sampled the novel odor more. (**C**) *'When' task*: Both WT groups and KOs given OXT (KO-OXT) spent more time sampling the earlier (Older) presented odor vs. the later (Recent) presented odor, whereas the KO-SAL group sampled both odors similarly. (**D**) *'Where' task*: Sampling times for the KO-SAL group were comparable between the switched and stationary odor pairs, while both WT groups and the KO-OXT group sampled the switched pair for a greater amount of time. *Statistics*: 2-tailed paired t-test: n.s. $p>0.05$, *$p<0.05$, **$p<0.01$, ***$p<0.001$, ****$p<0.0001$. Detailed statistical analyses are provided in supplementary table 3. Data presented as mean ± SEM.

### Figure S3

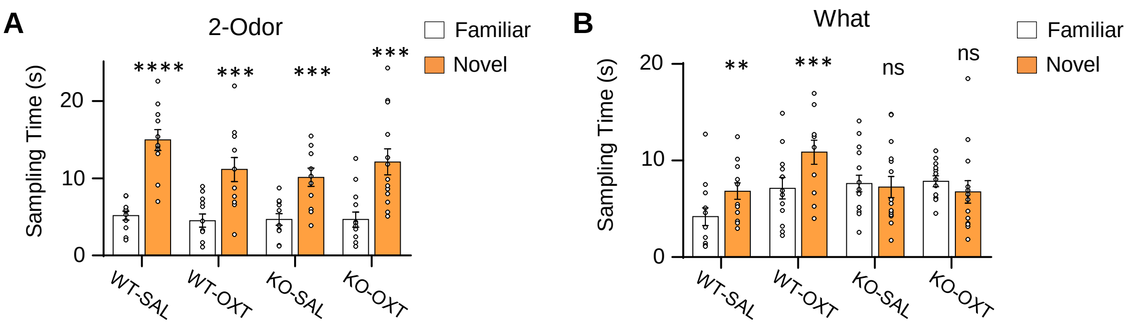


**Figure S3: Sampling times for 2-odor discrimination and episodic 'what' task in adolescence treated WT and KO groups.** (**A,B**) Sampling times (seconds) were used to calculate the discrimination indices presented in Fig. 5B (2-odor) and Fig. 5C ('What') for P30-P36 treated mice. (**A**) *2-Odor*: All four groups exhibited greater sampling times for the novel odor vs. the familiar odor. (**B**) *'What' task:* Both WT groups sampled the novel odor for greater time as compared to the familiar odor. By contrast, both the SAL-treated and OXT-treated KO groups sampled the two odors for similar amounts of time. *Statistics*: 2-tailed paired t-test: n.s. $p>0.05$, $^{**}p<0.01$, $^{***}p<0.001$. Detailed statistical analyses are provided in supplementary table 3. Data presented as mean ± SEM.
